# A difference in larval mosquito size allows a biocontrol agent to target the invasive species

**DOI:** 10.1101/2023.03.21.533626

**Authors:** Marie C. Russell

## Abstract

As the global temperature rises in the coming decades, *Aedes albopictus* is expected to invade and establish in South East England, where *Culex pipiens* is currently the most common native mosquito species. Biocontrol measures that use local cyclopoid copepods against *Ae. albopictus* may be compromised if the copepods prefer alternate *Cx. pipiens* prey. In this study, we assessed the predation efficiency of *Megacyclops viridis* copepods against French *Ae. albopictus* larvae and larvae that hatched from egg rafts of *Cx. pipiens* collected in South East England. The experiments were conducted at 15°C and 25°C, which are representative of present and future summer temperatures in South East England. *Ae. albopictus* larvae that survived the course of the experiment in the predator-absent controls were significantly smaller than *Cx. pipiens* larvae that survived in the absence of predation. The background mortality of *Cx. pipiens* larvae increased with the ten-degree increase in temperature, and the smaller size of surviving *Cx. pipiens* larvae at 25°C, relative to survivors at 15°C, suggests that larger *Cx. pipiens* larvae were more likely to die at the higher temperature setting. Across all experimental treatments, the ratio of copepod body length to mean prey length, based on larval lengths of survivors from the corresponding predator-absent controls, was a significant predictor of predation efficiency. Adding temperature setting to the predation efficiency model as a predictor did not improve model fit. Within the mixed prey treatments, the predation efficiency of *M. viridis* was 34.5 percentage points higher against *Ae. albopictus* prey than against *Cx. pipiens* prey. The higher predation efficiency that *M. viridis* exhibited against invasive *Ae. albopictus* prey, likely due to the smaller size of these larvae, supports the future use of *M. viridis* as a biocontrol agent in the UK.

## Introduction

Invasive *Ae. albopictus* mosquitoes lay desiccation-resistant eggs in artificial containers, such as tires (Benedict et al., 2007, Eritja et al., 2005, Juliano and Lounibos, 2005, Lounibos, 2002, Medlock et al., 2006), and are likely to establish in South East England within the next three to six decades (Kraemer et al., 2019, Metelmann et al., 2019, Proestos et al., 2015). *Cx. pipiens* is one of the most common species of mosquito in England (Golding et al., 2015), where it has been found in artificial containers, including tires (Chapman et al., 2017). A field study conducted in Italy found that 67% of all *Ae. albopictus* larvae shared their larval habitats with *Cx. pipiens* (Carrieri et al., 2003). In Italy, the efficiency of a cyclopoid copepod predator against newly-hatched *Cx. pipiens* larvae, 58.98%, was very similar to its predation efficiency of 54.99% against newly-hatched *Ae. albopictus* larvae (Veronesi et al., 2015). However, cyclopoid copepod predation efficiency among mixed larval prey populations has not been assessed using wild-caught *Cx. pipiens* from South East England and *Ae. albopictus* that originate from the temperate zone.

Other predator species, such as *Corethrella appendiculata*, have been shown to preferentially consume invasive *Ae. albopictus* larvae in the presence of native mosquito larvae (Griswold and Lounibos, 2006). Cyclopoid copepods feed on live prey by using mechanoreception to detect hydrodynamic disturbances (Awasthi et al., 2012, Roche, 1990). *Aedes* larvae have previously displayed higher motility than *Culex* larvae (Dieng et al., 2003, Kesavaraju et al., 2011), and higher larval activity has been associated with greater vulnerability to predation (Grill and Juliano, 1996). These findings are consistent with studies that have shown a preference for *Aedes* prey, rather than *Culex*, among cyclopoid copepod predators (Dieng et al., 2003, Soumare and Cilek, 2011, Pauly et al., 2022). However, the body size of prey organisms is also an important determinant of vulnerability to predation; cyclopoid copepods have been shown to prefer smaller species of rotifer prey and have caused greater mortality among fish larvae of smaller body lengths (Kumar et al., 2012, Lapesa et al., 2002).

In this study, we assess the predation efficiency of *Megacyclops viridis* copepods from Surrey, UK, which have previously been recommended as suitable biocontrol agents against invasive mosquitoes (Russell et al., 2021). The prey organisms include progeny of wild-caught *Cx. pipiens* from Berkshire, UK and *Ae. albopictus* from Montpellier, France. To reflect the range of both current summer temperatures and temperatures expected in the next three to six decades, we assessed predation efficiency at both 15 and 25°C. Based on previous studies, we do not expect temperature to have a significant effect on predation (Novich et al., 2014, Russell et al., 2021).

## Materials and Methods

### Collection of *Cx. pipiens* egg rafts from gravid females

Adult gravid female *Culex* mosquitoes were collected on the 24^th^ of June and the 13^th^ of July of 2019 from Ascot, Berkshire, UK near Silwood Lake (N 51°24.876’, W 0°39.045’) using a CDC gravid trap from the John W. Hock company (Model 1712). The trap was set on the preceding evening with 4L of hay infusion as the oviposition attractant. Once captured, each gravid *Culex* female was aspirated into a 25 cm x 25 cm x 25 cm cage and provided with 10% sucrose solution. A black plastic cup, containing 100 mL spring water, 15 mg spirulina,

33 mg Tetramin® fish food, 33 mg Russell Rabbit Tasty Nugget® rabbit pellets, and 33 mg powdered liver, was also provided for oviposition. The gravid *Culex* females were held in individual cages at 20 ± 1°C in a 12:12 light/dark cycle with 70% relative humidity until they laid an egg raft. After oviposition, the adult females were frozen at -20°C and identified by morphology as *Cx. pipiens/torrentium*. Because it is impossible to distinguish between *Cx. pipiens* and *Cx. torrentium* by morphology, each female was stored at -80°C, and eventually a portion of the 3’ region of the mitochondrial COI gene was sequenced in order to make a genetic identification (Hesson et al., 2010).

### Temperate *Ae. albopictus* colony care

A colony of *Ae. albopictus* mosquitoes (original collection Montpellier, France 2016, obtained through Infravec2) was maintained at 27 ± 1°C, 70% relative humidity, and a 12:12 light/dark cycle. Larvae were fed fish food (Cichlid Gold Hikari®, Japan), and adults were given 10% sucrose solution and horse blood (First Link Ltd, UK) administered through a membrane feeding system (Hemotek®, Blackburn, UK).

### *M. viridis* copepod cultures

Adult gravid female cyclopoid copepods were collected in May of 2019 from the edge of Longside Lake in Egham, Surrey, UK (N 51° 24.298’, W 0° 32.599’), and separate cultures were started from each gravid female. Each culture was kept in a 3 L container of spring water (Highland Spring, UK) at a 12:12 light/dark cycle, and 20 ± 1°C. *Chilomonas paramecium* and *Paramecium caudatum* (Sciento, UK) were provided *ad libitum* as food. Adult copepods were identified as *Megacyclops viridis* (Jurine, 1820) by Dr. Maria Hołyńska from the Museum and Institute of Zoology in Warsaw, Poland.

### Experimental design

Cages holding gravid *Culex* females were checked each day at noon for egg rafts. When egg rafts were found, plans were made to hatch the corresponding number of *Ae. albopictus* eggs beginning at midnight, 36 hours after finding the *Culex* egg raft (Fig 1). This schedule allowed there to be enough newly-hatched larvae of each genus by noon, 48 hours after the new *Culex* egg rafts were first observed.

**Fig 1.**
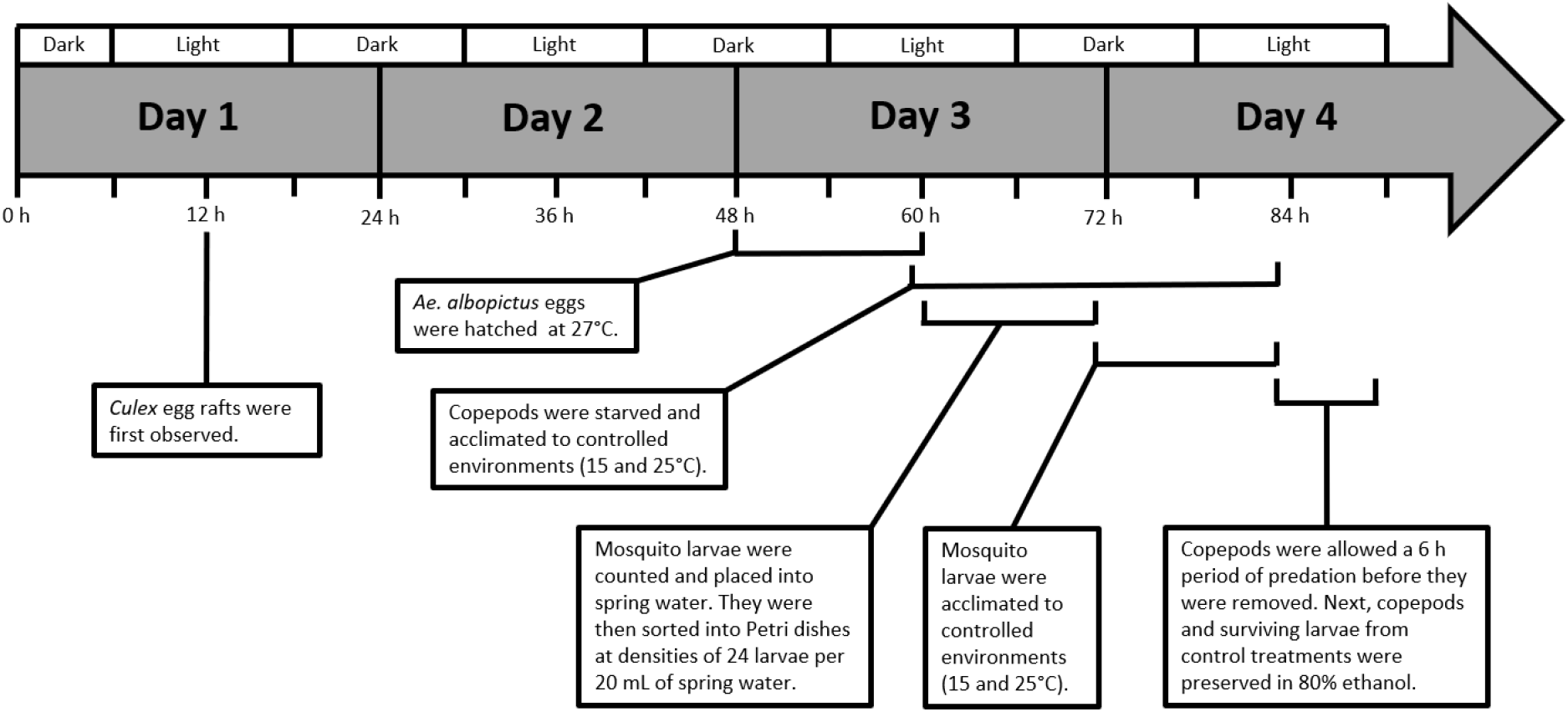
Schedule for predation efficiency experiment with native and invasive mosquito prey.

Adult non-gravid female *M. viridis* were each placed in a Petri dish holding 20 mL spring water for a 24 h period of starvation and acclimation to two different temperature settings: 15 ± 1 and 25 ± 1 °C, both at a 12:12 light/dark cycle. Mosquito larvae were counted and divided into Petri dishes so that one third of the Petri dishes held 24 *Culex* larvae, one third held 12 *Culex* larvae and 12 *Ae. albopictus* larvae, and one third held 24 *Ae. albopictus* larvae (Fig 2); half of these Petri dishes were acclimated to 15 ± 1°C, and the other half were acclimated to 25 ± 1 °C. Copepods were introduced to larvae in half of the Petri dishes for a 6 h period of predation. After the predation period, copepods were removed and the numbers of surviving larvae were counted and recorded. Copepods and surviving mosquito larvae from the control treatments were stored in 80% ethanol. The experimental schedule is displayed in Fig 1.

**Fig 2.**
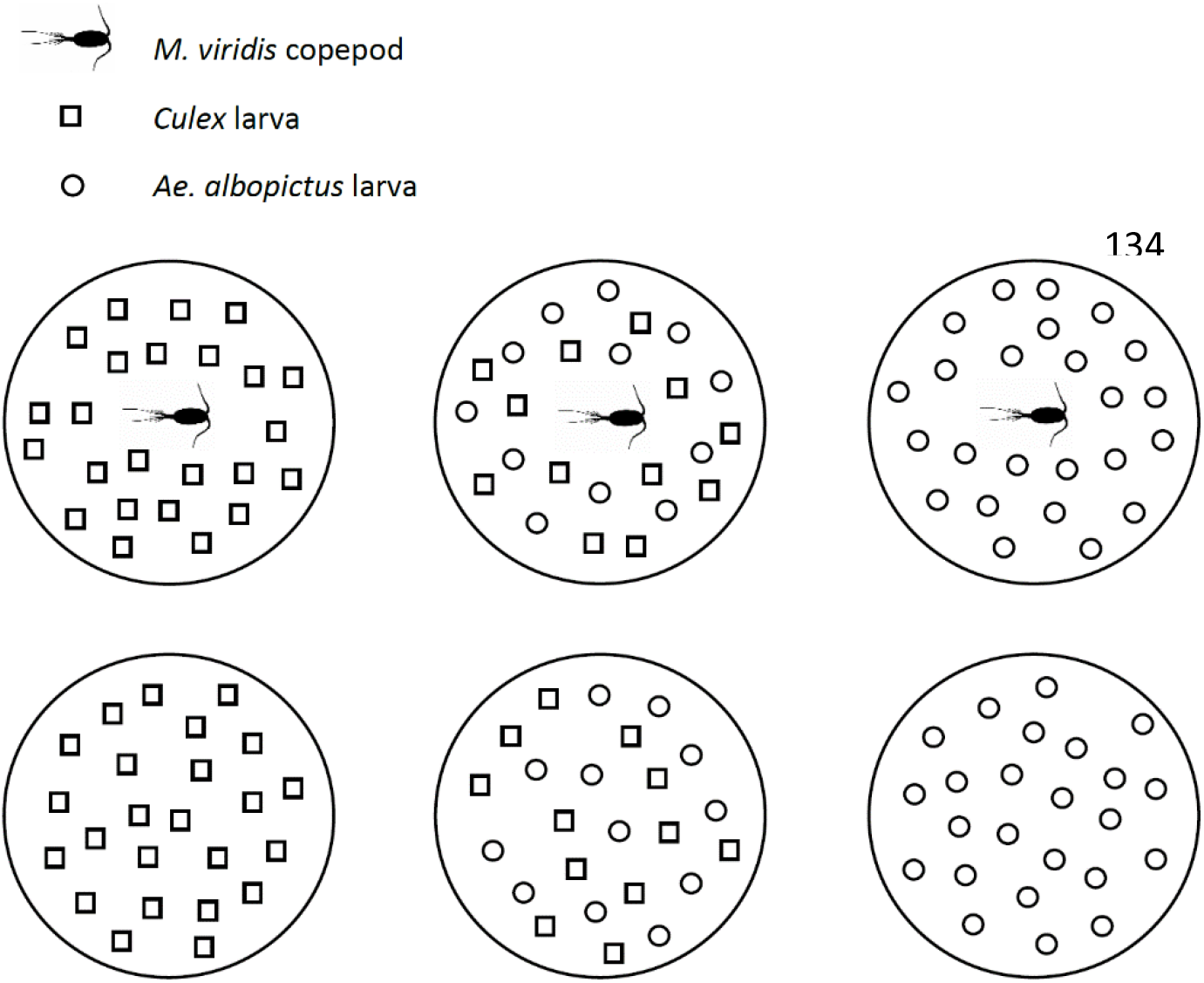
Experimental design repeated 16 times for each combination of *Culex* egg raft (8) and temperature (2). (Source of copepod silhouette accessed August 2019: http://phylopic.org/name/f79c5b86-7b73-468a-9ad4-995646398f99)

### Copepod and mosquito larva length measurements

The body lengths of all copepods, from the front of the cephalosome to the end of the last urosomite, were measured to the nearest tenth of a mm using a light microscope with an ocular scale bar. The body lengths of all surviving mosquito larvae in the control treatments were measured to the nearest hundredth of a mm, from the front of the head to the end of the saddle, using a ZEISS AxioCam microscope camera and AxioVision software (4.6.3).

### Statistical analyses

All analyses were conducted in R version 4.0.2 (R Core Team, 2020). Pearson’s chi-squared tests were used to determine if temperature affected the survival of mosquito larvae of each genus in the controls. A linear regression model was fitted to test mosquito genus and temperature as predictors of larval length. An additional linear regression model was fitted to test egg raft and temperature as predictors of *Culex* larval length.

Predation efficiency was calculated according to Abbott’s formula (Abbott, 1987, Baldacchino et al., 2017):

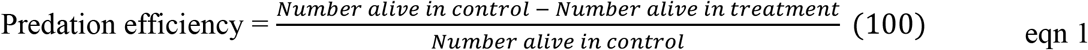

Copepod body mass (mg) was also calculated using an equation from previous studies (Alcaraz and Strickler, 1988, Klekowski and Shushkina, 1966, Novich et al., 2014):

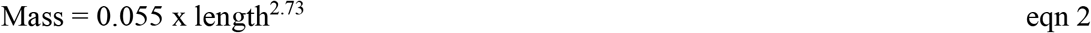

The mean prey length for each replicate (n = 48) was calculated based on the lengths of the surviving larvae in the controls. The predator-prey size ratio was then calculated by dividing the copepod length by the mean prey length. The size ratio values were restricted to two significant figures because the copepod lengths had two significant figures.

A linear regression model was fitted to test copepod length, copepod body mass, or copepod-to-larva size ratio as predictors of predation efficiency. Temperature, as well as an interaction between temperature and size ratio, were also tested as predictors. Replicates that had greater than 75% larval mortality in the control (n = 2) were excluded from predation efficiency analyses. In addition, one replicate where the copepod died during the period of predation was excluded.

Within each mixed prey replicate (n = 16), predation efficiency was also calculated as two separate values, each specific to a different genus of mosquito prey. One replicate was excluded because zero *Culex* larvae survived in the control. To assess if the *M. viridis* copepods preferred one type of prey to another, a paired t-test was used to determine if the differences between mosquito genus-specific predation efficiencies within each replicate (n = 15) were significantly different than zero.

All data will be made accessible from the Dryad Digital Repository.

## Results

All eight *Culex* females that corresponded to the eight egg rafts used in the experiment were identified as *Cx. pipiens* based on morphology and the sequence of 830 base pairs in the 3’ region of the mitochondrial COI gene (Hesson et al., 2010).

The probability of survival among *Ae. albopictus* larvae was 86.8% at 15°C and 86.1% at 25°C, and did not differ significantly based on temperature (p-value = 0.8076). However, among *Cx. pipiens* larvae, the probability of survival was significantly lower at 25°C, where 48.6% survived, as opposed to 93.8% at 15°C (p-value < 0.0001). The results of a Welch two sample t-test showed that surviving *Cx. pipiens* larvae were significantly larger than surviving *Ae. albopictus* larvae (mean ± sd: *Cx. pipiens* = 1.64 ± 0.18 mm, *Ae. albopictus* = 1.36 ± 0.13 mm; p-value = <0.0001).

The larval length model that only had mosquito species as a predictor had a similar AIC value (difference of less than 2) to the model that tested for interaction between mosquito species and temperature (Table S1). The results of the full model (Table 1) show that the interaction term was significant, indicating that the relationship between larval length and temperature was modified by mosquito species (Fig. 3).

**Table 1.** Linear regression of the full surviving larval length model (n = 908).

| Parameter | Estimate | Standard Error | p-value | Adjusted R<sup>2</sup> |
| --- | --- | --- | --- | --- |
| Intercept | 1.338 | 0.028 | $<0.0001$ | 0.458 |
| Temperature (ref = 15°C) | 0.001 | 0.001 | 0.5158 |  |
| Mosquito species (*Ae. albopictus* = 0, *Cx. pipiens* = 1) | 0.398 | 0.042 | $<0.0001$ | |
| Temperature x Mosquito species | -0.006 | 0.002 | 0.0035 |  |

**Fig 3.**
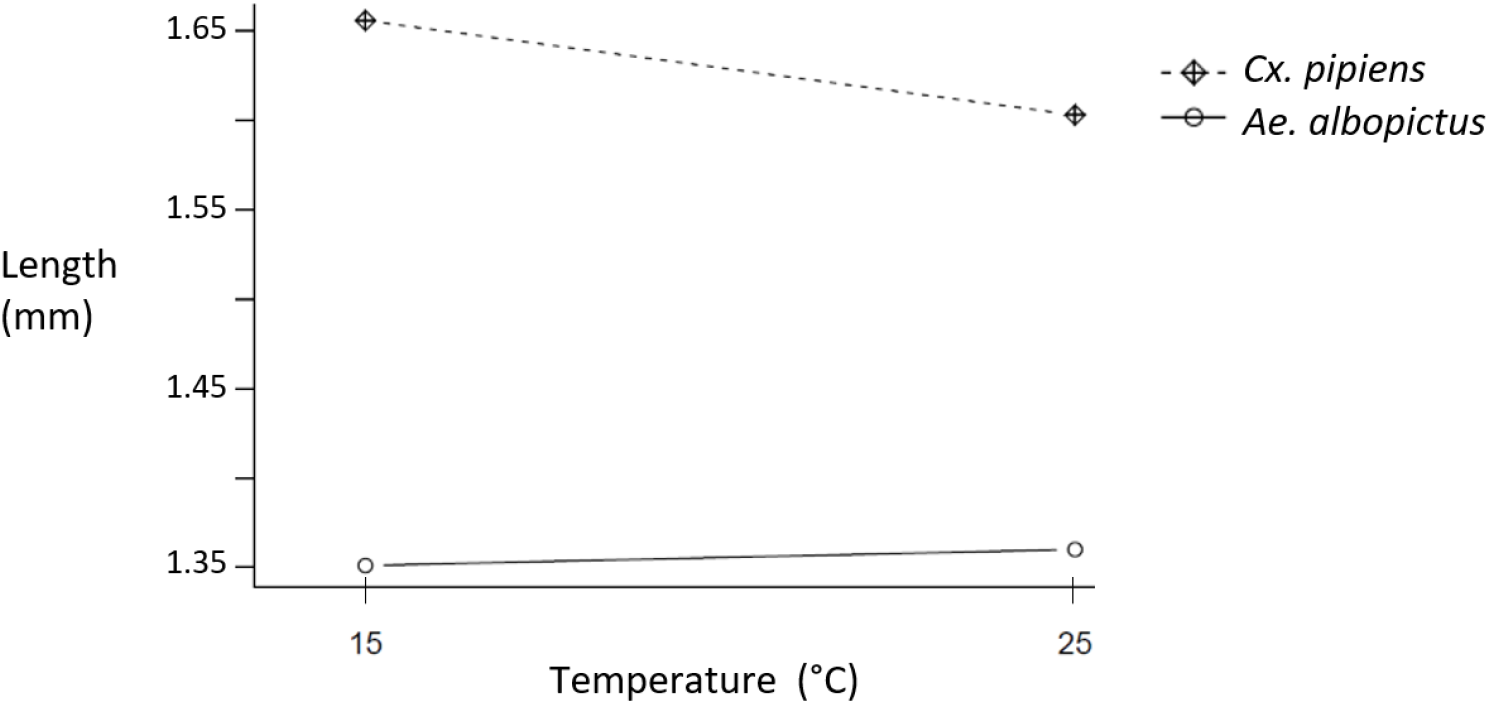
Interaction plot of relationship between surviving larval length and temperature, modified by mosquito species.

Further investigation of the larval lengths among only *Cx. pipiens* mosquitoes showed that the linear model with both egg raft and temperature as predictors had a similar AIC value (difference of less than 2) to the model that only included egg raft (Table S2). The results of the model that included temperature show that *Cx. pipiens* larval lengths significantly decrease with a 10°C increase in temperature, after controlling for natural variation in length due to egg raft effects (Table S3).

*M. viridis* lengths ranged from 1.5 to 2.5 mm (mean = 1.86, sd = 0.24). The predation efficiency model of best fit, based on AIC value, was the model that only included predator-prey size ratio as a predictor (Table S4). Adding temperature predictors to the model did not improve model fit (Table S4). Predator-prey size ratio was a significant predictor of predation efficiency (p-value = 0.0276), and for each unit increase in size ratio, predation efficiency increased by approximately 32% (Table 2, Fig. 4).

**Table 2.** Linear regression of predation efficiency by predator-prey size ratio (n = 45).

| Parameter | Estimate | Standard Error | p-value | Adjusted R<sup>2</sup> |
| --- | --- | --- | --- | --- |
| Intercept | -19.56 | 17.92 | 0.2810 | 0.087 |
| Predator-prey size ratio | 31.64 | 13.88 | 0.0276 |  |

**Fig 4.**
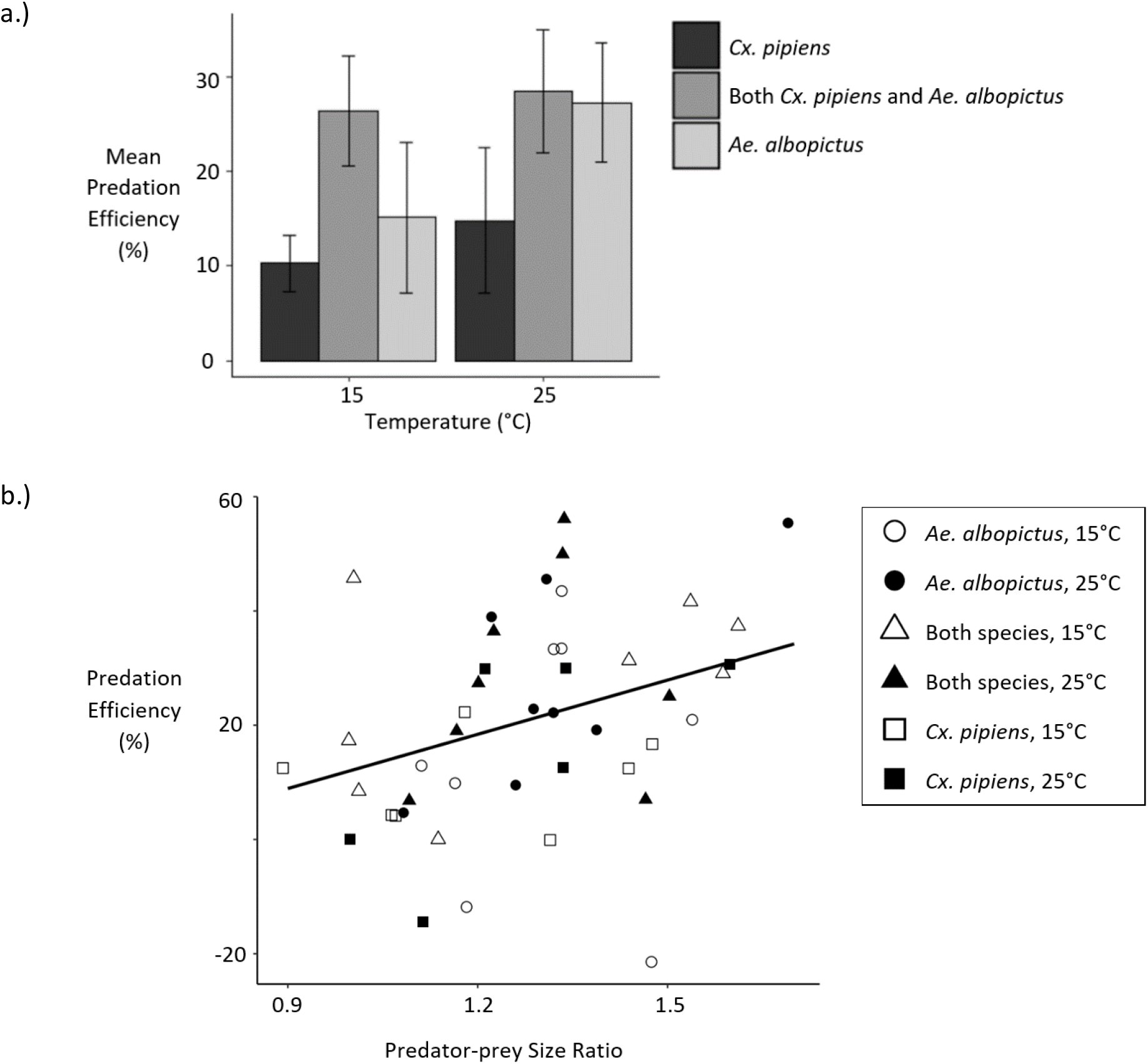
a.) Bar chart of predation efficiencies by temperature and prey species composition. b.) Scatter plot of predation efficiency by predator-prey size ratio.

The difference in prey species-specific predation efficiencies within each replicate that contained both *Cx. pipiens* and *Ae. albopictus* (n = 15) was significantly different than zero (p-value = 0.0107). On average, the *Ae. albopictus* predation efficiency was 34.5, 95% CI: (9.39, 59.70), percentage points higher than the *Cx. pipiens* predation efficiency (Fig. 5).

**Fig 5.**
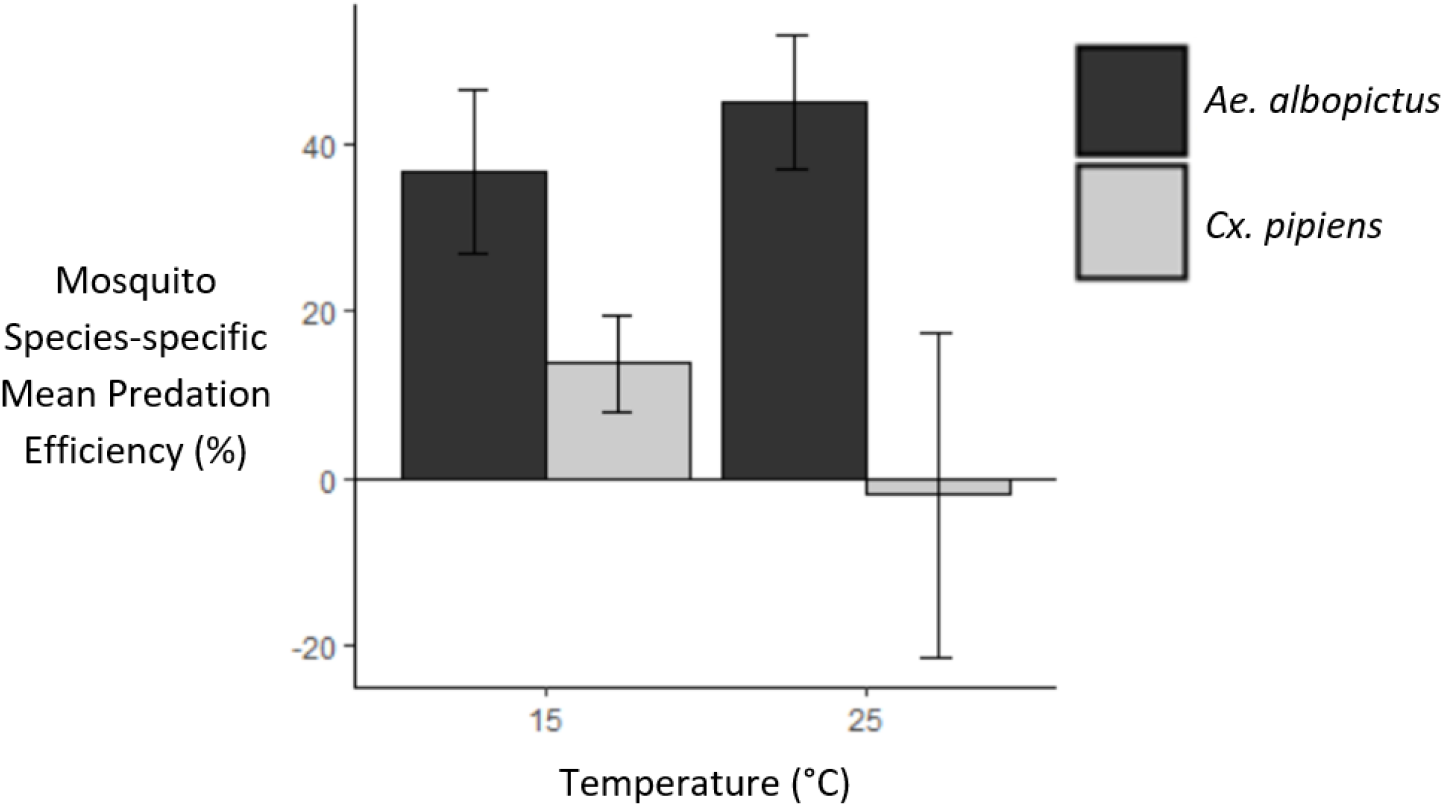
Mosquito species-specific predation efficiencies in treatments that contained both *Cx. pipiens* and *Ae. albopictus* larvae.

## Discussion

The survival of newly-hatched *Cx. pipiens* larvae was significantly lower at 25°C than at 15°C. Similar negative impacts of maintenance at 25°C have previously been observed for wild-caught UK mosquitoes of a different species, *Ae. detritus* (Lumley et al., 2018). Because the schedule of the experiment (Fig. 1) was too short to allow for temperature to strongly affect development, the inverse relationship between temperature and size observed among the *Cx. pipiens* larvae (Fig. 3, Table S3) was most likely mediated by higher mortality at 25°C. The reduction in size of surviving *Cx. pipiens* larvae at 25°C (Fig. 3, Table S3) is consistent with previous work showing that an increase in temperature can lead to a decrease in size among aquatic organisms (Daufresne et al., 2009).

Based on these results, colonies of *Cx. pipiens* that were maintained at 25°C (Hernandez-Triana et al., 2018, Manley et al., 2015) produced mosquitoes that were smaller than those found in natural UK populations. Lower colony maintenance temperatures, such as 22°C used for *Cx. pipiens* from Germany (Jourdan et al., 2016), or 23°C used for *Cx. pipiens* from the Netherlands (Manley et al., 2015), should also be used for *Cx. pipiens* from the UK. The low survival of *Cx. pipiens* larvae at 25°C also suggests that UK *Cx. pipiens* and *Ae. albopictus* might not share larval habitats as frequently as they have in regions at lower latitudes, such as Italy (Carrieri et al., 2003). Because *Cx. pipiens* larvae are known to inhabit both large permanent habitats (Amini et al., 2020, Lühken et al., 2015, Vinogradova, 2000), as well as container environments (Nikookar et al., 2017, Sulesco et al., 2015, Townroe and Callaghan, 2014, Verna, 2015), climate change may result in *Cx. pipiens* using the larger aquatic habitats more frequently, where water temperatures tend to be lower.

Predator-prey size ratio was a significant predictor of predation efficiency across all treatments (Table 2, Fig. 4). Previous work has shown that patterns in prey size selection are based on maximising the energetic gain of the predator (Mittelbach, 1981). For copepods, the optimal predator-to-prey size ratio has been estimated at 18:1 (Hansen et al., 1994). Based on this size ratio, both *Cx. pipiens* and *Ae. albopictus* larvae are much larger than the optimal prey for *M. viridis*. Additionally, it is expected that *M. viridis* would prefer the smaller larvae. Within treatments that had both prey species, *Ae. albopictus* predation efficiency was about 34.5 percentage points higher than *Cx. pipiens* predation efficiency (Fig. 5). The preference of *M. viridis* for *Ae. albopictus* prey supports its use as a biocontrol agent against these invasive mosquitoes. While species-specific larval behaviour may have also contributed to the higher vulnerability of *Ae. albopictus* (Kesavaraju et al., 2011), our analyses confirm that its smaller size played an important role.

Because *Aedes* mosquitoes are the costliest invasive taxon worldwide (Diagne et al., 2021), it is important to devise multi-faceted “integrated vector control” plans to keep mosquito populations as low as possible (Lacey and Orr, 1994). Our findings show that the efficacy of *M. viridis* biocontrol agents against *Ae. albopictus* prey would not be hindered in the presence of native *Cx. pipiens* larvae. This information suggests that including scheduled applications of *M. viridis* copepods to artificial containers could be a beneficial component of vector control plans in South East England.

## Acknowledgements

This project benefited from the advice and guidance of Dr. Lauren J. Cator. In addition, Claudia Wyer contributed to the molecular techniques that were employed for identifying *Culex pipiens* mosquitoes to species. Resources used during the experiments were funded by the European Union’s Horizon 2020 research and innovation program under grant agreement No 731060 (Infravec2). The work was primarily funded by a President’s PhD Scholarship from Imperial College London awarded to Marie C. Russell.

## Supplemental Material

**Table S1.** Surviving larval length model selection, ranked by AIC.

| Parameters | Degrees of Freedom | Adjusted R<sup>2</sup> | AIC |
| --- | --- | --- | --- |
| Mosquito species | 906 | 0.453 | -813.16 |
| Mosquito species, Temperature (ref = 15°C), Mosquito species x Temperature | 904 | 0.458 | -812.45 |
| Mosquito species, Temperature | 905 | 0.454 | -810.89 |

**Table S2.**
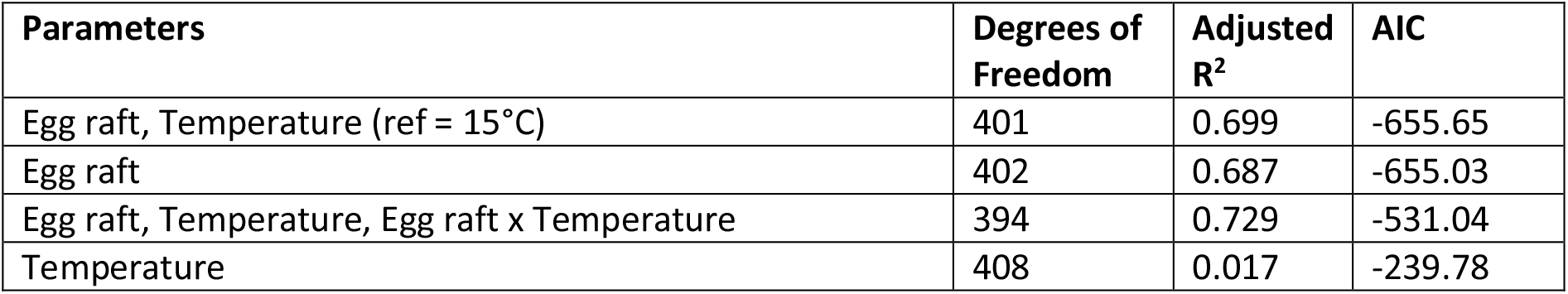
Surviving *Cx. pipiens* larval length model selection, ranked by AIC.

**Table S3.** Linear regression of surviving *Cx. pipiens* larval length by egg raft and temperature (n = 410).

| Parameter | Estimate | Standard Error | p-value | Adjusted R<sup>2</sup> |
| --- | --- | --- | --- | --- |
| Intercept | 1.427 | 0.026 | <0.0001 | 0.699 |
| Temperature (ref = 15°C) | -0.005 | 0.001 | <0.0001 |  |
| Egg raft 2 (ref = Egg raft 1) | 0.139 | 0.021 | <0.0001 |  |
| Egg raft 3 (ref = Egg raft 1) | 0.235 | 0.023 | <0.0001 |  |
| Egg raft 4 (ref = Egg raft 1) | 0.262 | 0.020 | <0.0001 |  |
| Egg raft 5 (ref = Egg raft 1) | 0.457 | 0.022 | <0.0001 |  |
| Egg raft 6 (ref = Egg raft 1) | 0.522 | 0.022 | <0.0001 |  |
| Egg raft 7 (ref = Egg raft 1) | 0.410 | 0.019 | <0.0001 |  |
| Egg raft 8 (ref = Egg raft 1) | 0.288 | 0.020 | <0.0001 |  |

**Table S4.** Predation efficiency model selection, ranked by AIC.

| Parameters | Degrees of Freedom | Adjusted R<sup>2</sup> | AIC |
| --- | --- | --- | --- |
| Predator-prey size ratio | 43 | 0.087 | 385.32 |
| Copepod length | 43 | 0.022 | 388.40 |
| Copepod body mass | 43 | 0.008 | 389.06 |
| Predator-prey size ratio, Temperature | 42 | 0.094 | 388.92 |
| Predator-prey size ratio, Temperature, Predator-prey size ratio x Temperature | 41 | 0.100 | 394.53 |

